# A multimodal human-computer interaction dataset for neurocognitive user state evaluation

**DOI:** 10.1101/2025.09.03.673947

**Authors:** Sai Zhang, Xinyu Bai, Charles Hartley-O’Dwyer, Hugh Warren, Frederike Beyer, Valdas Noreika

## Abstract

We introduce the Simulated Environment for Neurocognitive State Evaluation (SENSE-42), a multimodal dataset collected during user interactions with desktop computers. It is designed for studying spontaneous fluctuations in the neurocognitive state related to the tonic alertness of computer users, with recordings from 42 participants over 2-hour sessions. Within a simulated desktop environment, participants performed real-world routine tasks, including application switching, file management, typing, and web browsing. High-resolution data were recorded across physiological (electroencephalography, electrocardiography, respiration) and subjective modalities of alertness. At five-minute intervals, alertness state was reported using seven questions, addressing sleepiness (Karolinska Sleepiness Scale), mental and temporal demand, perceived performance, effort and frustration (NASA Task Load Index), as well as attentiveness. Behavioural data included keyboard, mouse and webcam inputs. Demographic information, experience metrics, habits, and preferences of computer usage were collected. In addition, individual differences in sleep quality were evaluated using the Pittsburgh Sleep Quality Index and the Epworth Sleepiness Scale. The SENSE-42 dataset can contribute to future research in user state monitoring, behavioural analysis and physiological computing.

## Background & Summary

Understanding and optimising task performance and interaction quality in human-computer interaction (HCI) is essential for maintaining productivity, improving user experience and minimising error-related safety hazards ^1;2^, particularly when users are engaged in prolonged and cognitively demanding activities on the computer. The performance and engagement of computer users are governed by their underlying neurocognitive states, which encompass cognitive, affective, and physiological processes, including attention, sleepiness, mental workload, and emotional responses such as frustration and perceived effort ^3;4;5^. Monitoring spontaneous fluctuations in the neurocognitive state related to the tonic alertness of computer users can reflect these internal states, supporting sustained attention and stable task engagement over time ^6;7^. Understanding and evaluating tonic alertness in structured, task-driven HCI contexts is therefore critical for reducing frustration, increasing engagement, and enhancing overall satisfaction, which forms the focus of this work.

Various approaches have been developed to assess alertness, broadly categorized into subjective and objective measurements. Subjective assessments, such as the Karolinska Sleepiness Scale (KSS) ^8^, the Epworth Sleepiness Scale (ESS) ^5^, and the NASA Task Load Index (NASA-TLX) ^4^, provide insight into perceived cognitive states but suffer from challenges in reliability and temporal resolution^9^. Objective methods, ranging from task-based performance metrics^10^ to physiological measures such as electroencephalogram (EEG) and cardiac signals ^11;12^, offer greater temporal granularity but are often more intrusive or technically demanding. Subjective measures provide a common scale for rating perceived alertness, enabling comparability across individuals and contexts. Conversely, objective measures facilitate continuous, minimally disruptive monitoring, mitigating the inherent limitations of self-reported data such as recall bias^13^ or momentary introspective inaccuracy ^14^. Together, these approaches offer complementary strengths, enabling a more accurate and holistic modelling of internal user states.

Prior studies in HCI have shown substantial progress in linking user alertness with behavioural cues derived from keyboard usage, mouse movement, and eye-tracking data ^15;16;17^. Pimenta et al. investigated the monitoring of mental fatigue through keyboard and mouse interaction patterns, focusing on generic, task-independent metrics such as keydown time, error rate per key press, mouse acceleration, and double-click speed, collected during two sessions at the start and end of each day^15^. Yamada et al. examined the detection of mental fatigue using eye-tracking data before and after cognitively demanding tasks, reporting improved accuracy over earlier approaches ^16^. Natnithikarat et al. proposed a drowsiness detection method that integrates keyboard, mouse, and eye-tracking dynamics in a standalone application to simulate a working routine in the accounting division and perform automated quality checks on transactions ^17^. While these efforts have demonstrated promising associations, several limitations remain. Most existing datasets rely heavily on subjective self-report measures, lack multimodal integration, and are based on unscheduled or loosely structured tasks without strict timings or a fixed experimental schedule, limiting their intrinsic validity and reliability^18;19^. Moreover, few studies have focused on the continuous measurement of spontaneous fluctuations in user states, with most studies comparing only discrete time points such as before and after tasks or morning and evening effects ^20^. Furthermore, few studies incorporate synchronised physiological and behavioural data in real-world-like computer usage scenarios ^21^. The absence of reliable physiological data source from EEG, ECG or respiration limits the ability to link internal physiological states with external observable user behaviours, hindering the development of more accurate models and predictions of neurocognitive states across broader applications.

To address these gaps, we introduce a multimodal dataset named **S**imulated **E**nvironment for **N**eurocognitive **S**tate **E**valuation (SENSE-42) ^22^, designed to study early alertness state fluctuations during realistic computer interaction in a desktop operating system environment. To accommodate user preferences while maintaining a unified experimental design, we developed precise replicas of two major operating system interfaces (macOS^23^ and Windows ^24^) to facilitate the study of task-related behavioural responses. Both interface styles were presented to each participant during the session, allowing within-subject comparisons across familiar and unfamiliar environments. The dataset includes synchronised high-resolution neurocognitive responses in multiple modalities, including physiological, behavioural, and self-reported data, from 42 participants within a simulated desktop operating system. These responses were collected over continuous 2-hour sessions for each participant, providing rich measurements of spontaneous fluctuations in user states. Structured tasks were designed to emulate everyday computer use—file management, typing, browsing, and application switching—under tightly controlled conditions. Participants self-reported their alertness state at five-minute intervals using predefined scales, including the Karolinska Sleepiness Scale (KSS) ^8^, NASA Task Load Index (NASA-TLX)^4^, and attentiveness ratings ^25^. Physiological data were collected to profile internal user states from EEG, ECG and respiration signals, and behavioural data were recorded for keyboard and mouse dynamics. Additionally, trait-level measures such as the Epworth Sleepiness Scale (ESS) ^5^ and the Pittsburgh Sleep Quality Index (PSQI) ^26^ were collected to contextualise inter-individual variability in daytime sleepiness and nighttime sleep patterns.

The SENSE-42 dataset^22^ is collected in a time-locked experimental setting to replicate real-world computer use scenarios. It provides a structured resource for examining tonic alertness dynamics during human-computer interaction and may be used to explore relationships between various multimodal indicators of the user state and their personal traits. The dataset includes components relevant to research in areas such as cognitive state monitoring, user behaviour analysis, and physiological computing^27^.

## Methods

### Participants

A total of 42 participants were recruited in our study, consisting of 21 males and 21 females. The average age of participants is 26.8 ± 4.5 years, ranging from 18 to 40. Participants represented diverse ethnic backgrounds, education levels, and occupations. To preserve ecological validity, they were instructed to maintain their usual intake of substances that could influence alertness, such as caffeine or medication. The study was approved by the Queen Mary Ethics of Research Committee (Reference Number: PSY2025-32), and the procedures were designed in accordance with the Data Protection Privacy Notice of Queen Mary University of London ^28^. Detailed explanations of the study and its procedures were provided to the participants, and informed consent for data publication with optional data-sharing preferences was obtained from all participants. Participants received £30 for taking part in the study.

### Study Design

Data collection was organised from March 2025 to July 2025 in the Centre for Brain and Behaviour at Queen Mary University of London. The overall experiment design is shown in Fig. 1 (a). The study followed a single-group observational design, with all participants completing an identical task sequence in a fixed order. Additionally, a within-subject design was applied to evaluate user interactions across two conventional interface layouts within the simulated operating system (see Fig. 1 (b)). This dataset ^22^ includes all EEG, ECG, respiration, keyboard, and mouse data from the calibration stage to the end of the two-hour session. The experiment sessions were conducted individually, with participants selecting their preferred time slot from 11:00, 13:00, or 15:00. Participants’ gender and operating system layout preferences were balanced at a 50:50 ratio to reduce response biases, and only individuals proficient with English and desktop computers were included to ensure consistent task engagement. The experiment was conducted in a sound-insulated chamber, with the experimenter seated in a separate control room to minimise external influence. Environmental conditions were strictly controlled: ambient noise was maintained at approximately 52 dB, room temperature was kept at normal range (∼25°C), lighting was controlled at standard workplace levels (∼500 Lux), and *CO*_2_ concentration was kept below 800 ppm.

**Figure 1:**
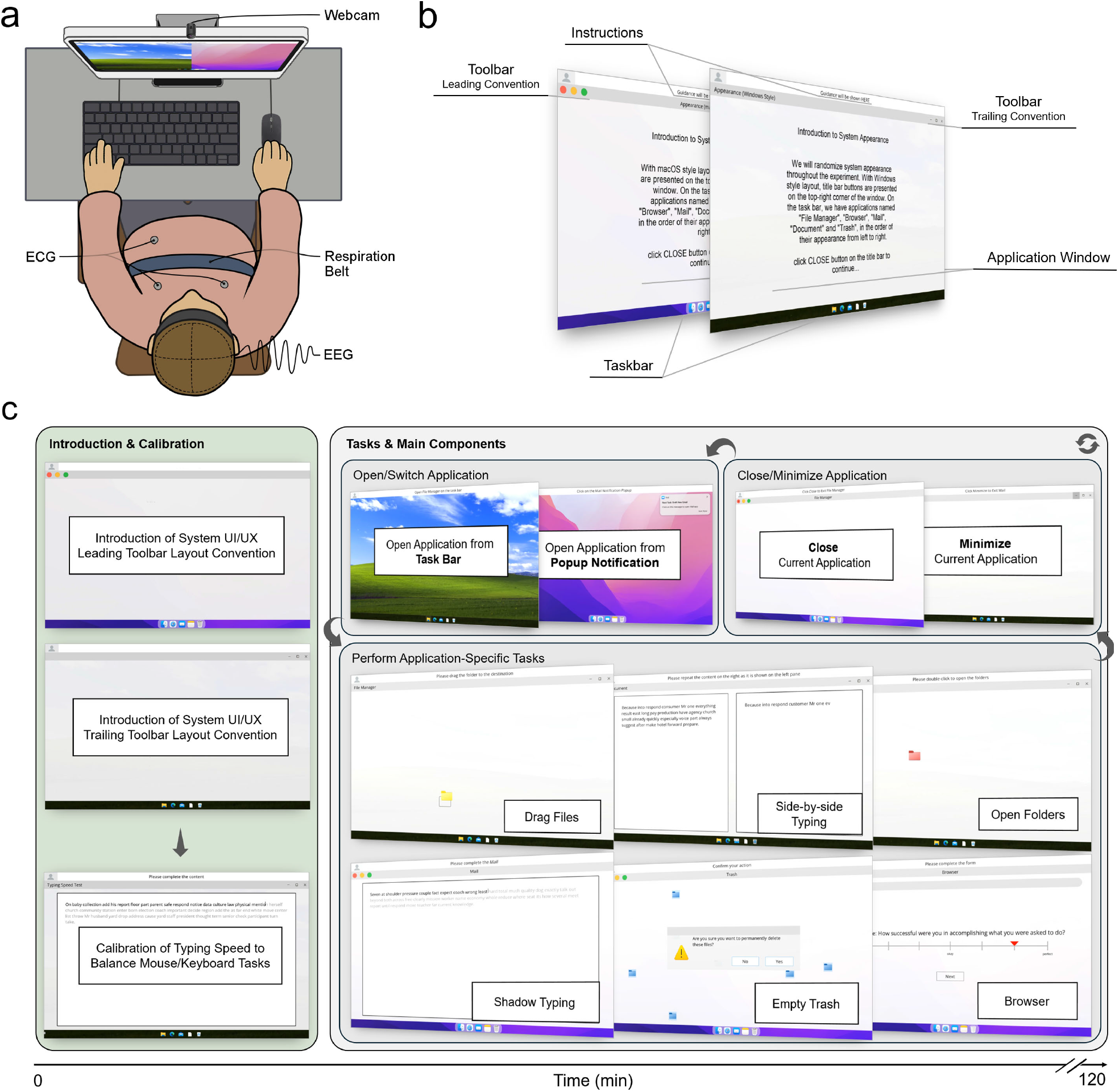
Experiment design. a) Top view of the experiment setup. Participants were asked to perform tasks on a computer following instructions at the top of the screen, with keyboard and mouse as the main input devices. EEG, ECG, respiration data, and webcam recordings were collected throughout the session. b) Illustration of software UI/UX design complying with two conventional operating system layouts. c) Details of experiment flow with screenshots. The duration of the experiment session is two hours.

### Experimental Design

The illustration of the overall experimental design is shown in Fig. 1 (c). The experiment began with an introduction and calibration phase in which participants were introduced to two conventional operating system layouts (toolbar items in leading position: macOS ^23^; toolbar items in trailing position: Windows ^24^) and their corresponding UI/UX elements. Typing speed was calibrated in the next block to ensure time balance between keyboard- and mouse-based tasks.

The main experiment lasted approximately two hours and consisted of a fixed set of individual or chained tasks within a fully controlled and simulated software environment. Tasks included: (1) opening or switching applications and responding to mail notifications; (2) performing application-specific operations such as dragging/opening folders, shadow typing or side-by-side typing, emptying the trash folder, or browsing webpages; and (3) closing or minimising the current application.

Participants were instructed to open designated applications either by clicking the corresponding icon on the taskbar located at the bottom of the screen or by clicking on the mail notifications appearing on the right side of the screen after a 5-second pause.

Once an application was launched, participants performed application-specific tasks within a continuous workflow. Mouse navigation tasks were confined within the central area of the application main window, occupying half of its width and height. Dragging folder tasks were repeated 20 times and opening folder tasks were repeated 30 times, with stimuli location randomised in each iteration. Typing tasks used word stimuli generated from a vocabulary list sampled according to word occurrence frequencies via the Faker library ^29^ and were designed to last approximately one minute per block. Participants performed two types of typing tasks: (1) shadow typing, where the target text was displayed in a lighter colour and participants typed over it, gradually replacing the reference as they typed; and (2) side-by-side typing, where the reference text remained visible on the left side of the screen while participants entered their response on the right. For file management, participants selected a set of files and confirmed their deletion via a pop-up dialogue. Web browsing tasks involved entering a typical URL generated by the Faker library ^29^ and completing the subject questionnaires embedded in a webpage every five minutes to assess their current state across seven dimensions: sleepiness (KSS) ^8^, mental demand, temporal demand, performance, effort, frustration (NASA-TLX)^4^, and attentiveness (assessed with momentary mind-wandering scales) ^25^.

Finally, participants were instructed to hide the application window by clicking the minimise or close button located on the top toolbar.

The order of tasks within each block was fixed to maintain a balance between mouse- and keyboard-driven interactions, while the system layout was randomly assigned between the two with equal probability and remained consistent throughout the same block. The questionnaires were presented in a non-intrusive manner, designed to minimally disrupt task engagement. The timing of questionnaire presentation was linked to the preceding task duration and controlled to appear approximately every five minutes. A screen recording of the experiment is provided in Supplementary Video S1.

Throughout the session, multimodal data were continuously recorded, including EEG, ECG, respiration, behavioural interactions via mouse & keyboard, and self-reported responses. Within the behavioural data, mouse and keyboard responses including time, duration and location of each key press were synchronised with the screen refresh at 144 Hz and recorded on each frame. The timing of all stimulus presentations is included in the dataset, with the start of each test block linked to EEG triggers. Setup of physiological equipment was completed concurrently with the administration of pre-study questionnaires. Experiment guidelines were offered at the start of the experiment, encouraging participants to work efficiently and minimise pause between blocks.

### Experimental Setup

The experiment software was implemented using PsychoPy v2024.2.3^30^ and run on a Dell Precision 3490 workstation (Windows 11 Pro 24H2^24^) equipped with an RTX 500 Ada GPU. The display was presented on a 144 Hz Dell G3223Q monitor, operating at the resolution of 1920×1080 with frame-to-frame refresh stability maintained below 1.5 ms. Cursor speed was set to the system’s medium setting. Participants interacted with a Dell SK-8175 keyboard and a Logitech M-U0017 optical mouse on a mousepad. A Dell WB5023 webcam operating at 1280x720 (50Hz) was used for video recordings where participant consent was provided. EEG data were recorded using a BioSemi ActiveTwo system with 32 electrodes arranged according to the extended 10–20 international placement system. Data were sampled at 1024 Hz, with electrode impedance maintained below 5 *k*Ω. EEG, ECG and Respiration acquisition was controlled via ActiView v8.0 running on a macOS 10.11.6 workstation. Environmental conditions were monitored using a Newentor C1 air quality sensor. The monitor and input devices were wired into the sound-attenuated chamber from the control room.

## Data Records

The SENSE-42 dataset ^22^ (DOI: 10.5281/zenodo.20328099) is organised into six top-level directories reflecting the different modalities collected: Questionnaires, Behavioural, ECG, EEG, Respiration, and Webcam. Each directory contains flattened data files labelled with anonymised participant IDs. 3-lead ECG signals (RA, LA, LL) ^31^ were extracted from the auxiliary channels of the EEG recordings and saved in 3-channel .fif format using the MNE-Python library^32^. Respiration waveforms were collected with a respiration belt around the chest, sampled at 32 Hz and stored in .wav format, processed via the RespInPeace package ^33^. EEG recordings are shared in the original BioSemi Data Format (BDF) and cleaned EEGLab Format (.set) following the post-processing pipeline in the technical validation step of EEG data. Webcam recordings are stored as individual MP4 files at 720p resolution and 50 FPS, organised by participant with one file per test block. The collected webcam data is suitable for capturing block-level behaviour rather than for precise temporal alignment in fine-grained gaze estimation or pupillometry. All participants agreed to share their webcam data for authorised research use. Table 1 summarised the released set of pre- and in-study behavioural questionnaires. For details on how these questions were presented to participants, refer to the archived pre-study questionnaire document and the video recording of the experiment available in the dataset files. Further documentation, including field notes, contact details and an overview, is provided on the dataset page.

**Table 1:** Released behavioural data from pre-study and in-study questionnaires.

| Name | Key Variables |
| --- | --- |
| **Pre-study Questionnaire** |  |
| Demographic | Age, gender, occupation, ethnicity, highest level of education |
| Personal Traits | Mental health conditions, substance use, handedness |
| Task Proficiency | Preferred language, computer usage proficiency, preferred computer setup, keyboard usage frequency |
| Pittsburgh Sleep Quality Index (PSQI)<sup>26</sup> | Sleep habits, overall sleep quality |
| Epworth Sleepiness Scale (ESS)<sup>5</sup> | Total score |
| **In-study Questionnaire** |  |
| Karolinska Sleepiness Scale (KSS)<sup>8</sup> | Sleepiness rated on a 9-point scale |
| NASA Task Load Index (NASA-TLX)<sup>4</sup> | Mental demand, temporal demand, performance, effort, frustration rated on a 7-point scale |
| Task Engagement<sup>25</sup> | Attentiveness rated on a 7-point scale |

## Data Availability

The SENSE-42 dataset can be downloaded with an approved access request via Zenodo ^22^ (DOI: 10.5281/zenodo.20328099).

## Technical Validation

### Behavioural Data

Participants completed an average of 34 ± 5 task blocks, with each block lasting approximately 227 ± 57 seconds. Within these blocks, 16 ± 4 trials were arranged in leading system toolbar layout and 18 ± 3 trials were in trailing system toolbar layout. The mean time spent on each task was 11.72 ± 9.50 seconds for opening the application, 28.12 ± 8.76 seconds for dragging folders, 23.60 ± 3.75 seconds for opening folders, 54.21 ± 18.85 seconds for shadow typing, 59.84 ± 26.61 seconds for side-by-side typing, 33.85 ± 18.48 seconds for web browsing, 3.44 ± 2.38 seconds for emptying the trash bin and 2.24 ± 1.63 seconds for closing or minimising the application. To validate task engagement and layout-specific behaviour, we present mouse navigation heatmaps separately for the two system layout conventions in Fig. 2 (a). Mouse locations are mainly distributed around the taskbar, toolbar and the centre of application window. Additionally, logarithm count of key-presses on the keyboard are visualised in Fig. 2 (b). The frequent appearance of letter E, T, A, O and rare occurrence of letter Z, J, Q, X are aligned with the previous study on the letter frequencies in the English vocabulary ^34^. These statistics support the validity and reliability of the recorded behavioural responses across participants.

**Figure 2:**
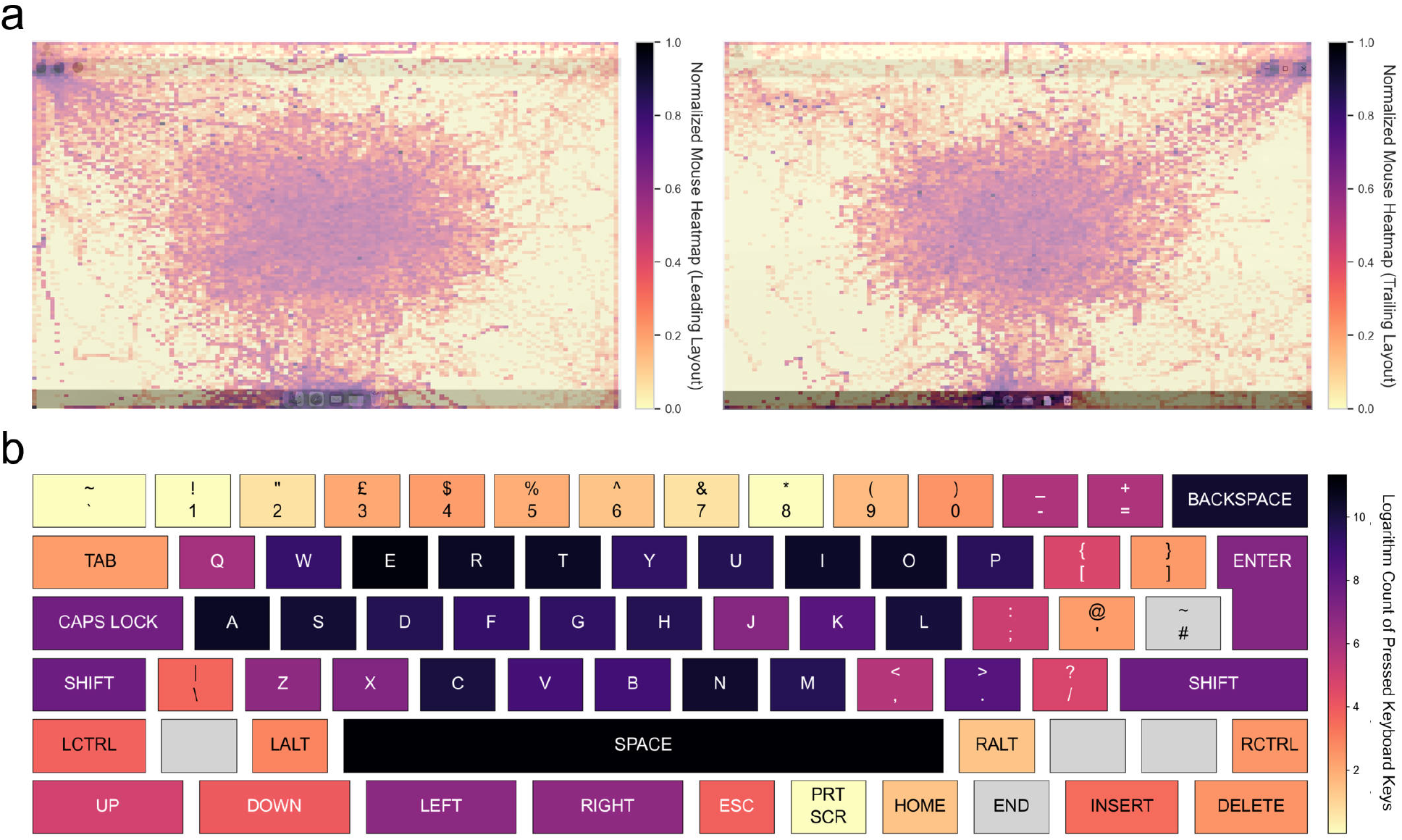
Behavioural data validation through keyboard and mouse interaction records. a) Normalised mouse location heatmap on the screen under leading and trailing system layouts. The central concentration in both heatmaps indicates frequent common task engagement, while peripheral activity reflects interactions with operating system components. b) Logarithm count of keyboard key presses, mapped onto a standard UK keyboard layout. Darker colours indicate more frequent hits, demonstrating consistent interaction patterns with frequently used function and character keys.

### Physiological Data

To validate the physiological data quality and consistency over time, we applied standardised preprocessing and analysis pipelines across EEG, ECG, and respiration signals. We checked critical quality indicators for each physiological sensor, including the resting-state EEG power spectral density (PSD), event-related potential (ERP) associated with the visual onset of folder stimuli, distribution of cardiac timing metrics and distribution of respiration timing metrics.

EEG data were processed using the Automagic toolbox ^35^ in MATLAB R2023a^36^, which facilitated automatic detection and removal of muscle and eye movement artifacts, identification and rejection of noisy channels, followed by spherical spline interpolation for identified bad channels and rejection of noisy segments. After preprocessing, we further computed the power spectral density for Cz using the Welch method ^37^. The absolute power of each electrode was transformed into the logarithmic scale (1*dB* = 10 log(*µV*^2^)) to represent the relative signal strength. The averaged power spectrum at Cz channel for all participants is shown in Fig. 3 (a), and the power spectral density plots for each participant are presented in Supplementary Material S2. The spectrum fits the criteria of good data quality with a smooth decline towards higher frequency and a slight increase around the alpha band power ^38^. The averaged event-related potential time-locked to the presentation of the computer home screen at Fz channel is presented in Fig. 3 (b). Extracted epochs were de-trended and baseline-corrected using the interval -100 ms to 0 ms. The averaged waveform shows a clear negative deflection (N100) peaking at approximately 120 ms, followed by a positive deflection (P200) around 180 ms after stimulus onset. These components are consistent with well-established visual ERP responses, where the N100 is associated with early sensory processing and the P200 with subsequent perceptual evaluation and attention allocation ^38^. Standard deviations are marked in gray in both figures.

**Figure 3:**
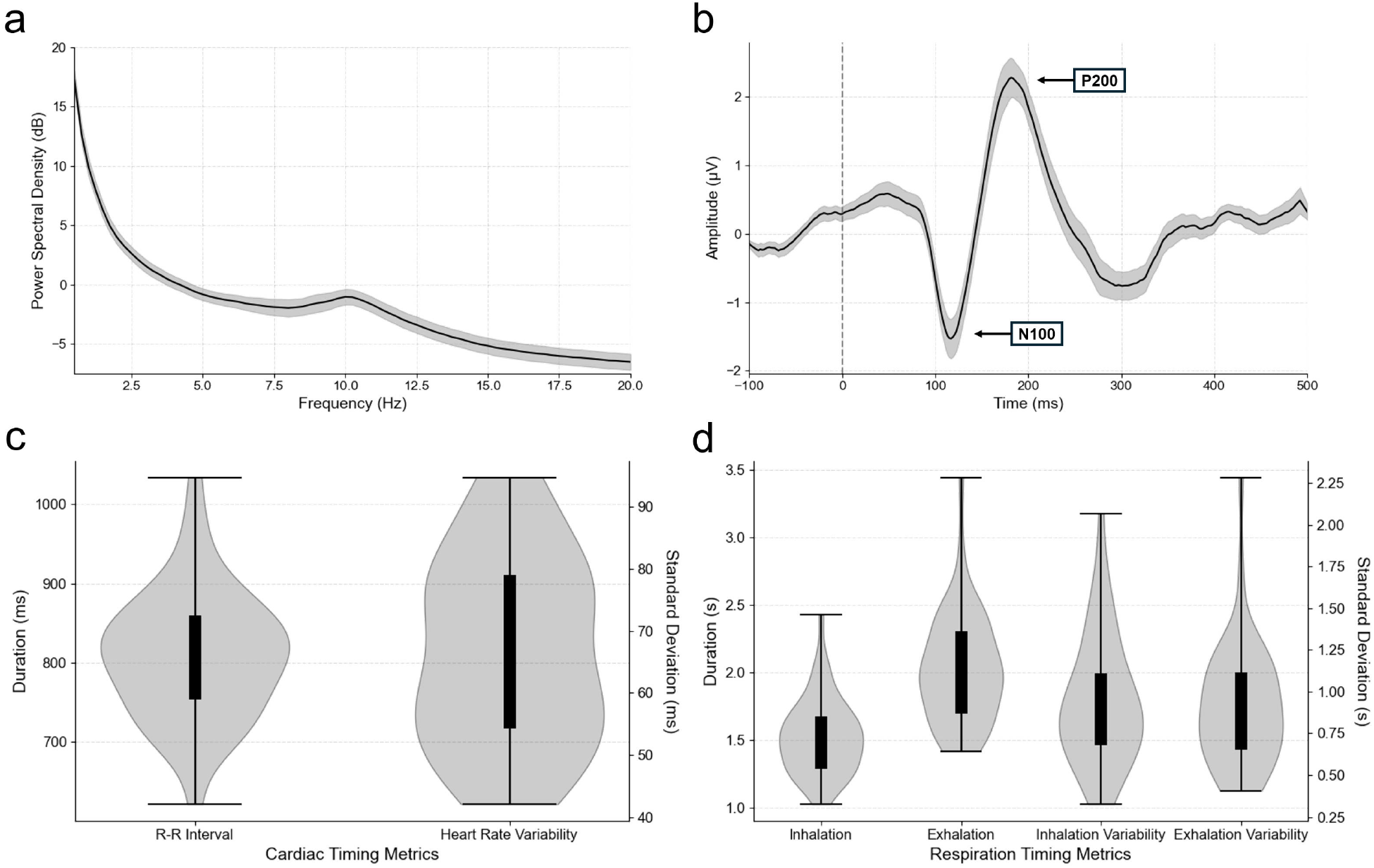
Physiological data validation through EEG, ECG, and respiration signals. a) Power spectral density (PSD) of Cz channel within 0-20 Hz, averaged across participants with standard deviations shown in gray. b) Event-related Potential (ERP) time-locked to visual stimulus onset at Fz channel, averaged across participants with standard deviations in gray. The vertical dashed line at 0 s denotes the appearance of the visual cue (computer home screen). c) Cardiac timing metrics for R-R interval and heart rate variability derived from 3-lead ECG recordings. d) Respiration timing metrics with inhalation and exhalation patterns.

For cardiac signals, we extracted events from a derived 3-lead ECG time series and selected lead II signals calculated from the difference between the left leg (LL) and right arm (RA) to be processed by the MNE-Python^32^. Only valid heartbeats within the physiological range (50–220 bpm) were retained. We calculated mean R–R intervals and heart rate variability (HRV) from Standard Deviation of N-N intervals (SDNN) ^39^ as indicators of autonomic activity (as in Fig. 3 (c)). The observed ranges for R–R intervals and HRV are within established short-term normative values ^40^, indicating typical cardiac function in the recorded data.

Respiration signals were processed using the RespInPeace package ^33^, which applied low-pass filtering for noise reduction, baseline estimation to account for temporal drifting, and keypoint identification for inhalation and exhalation. Based on the RespInPeace package, we derived breathing-related duration and variability metrics for comparison in Fig. 3 (d). The results are consistent with expected physiological ranges for healthy adults ^41^, indicating that the recorded breathing patterns were within normal limits.

### Self-reported Data

Participants self-reported their attentiveness, mental demand, temporal demand, performance, effort, frustration, and sleepiness during the study, as shown in Fig. 4. Sleepiness was assessed on a 9-point scale using the Karolinska Sleepiness Scale (KSS)^8^, while the other six dimensions were rated on a 7-point scale adapted from the NASA Task Load Index (NASA-TLX) ^4^ and momentary mind-wandering assessments ^25^. The self-assessment blocks were arranged every 299 ± 43 seconds, consistent with the intended five-minute schedule.

**Figure 4:**
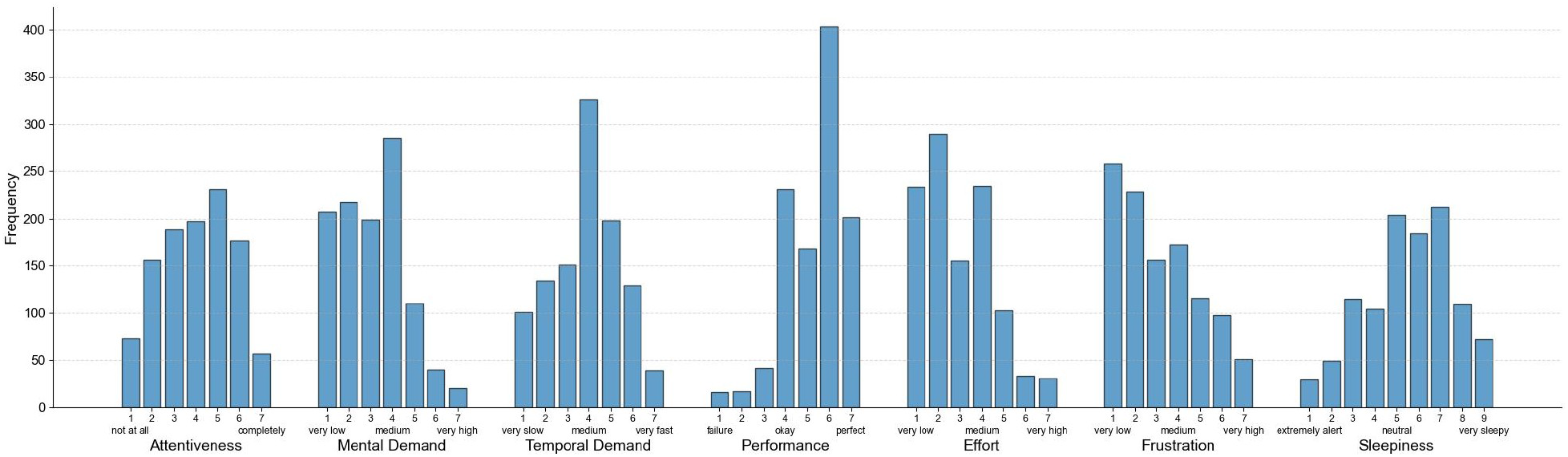
Distribution of self-reported ratings for attentiveness, mental demand, temporal demand, performance, effort, frustration, and sleepiness across all trials. Ratings for all metrics except sleepiness were measured using a 7-point scale adapted from NASA-TLX ^4^ and momentary mind-wandering assessments ^25^, while sleepiness was assessed using the 9-point Karolinska Sleepiness Scale (KSS) ^8^.

### Multi-modality Sensor Synchronisation

To validate the synchronisation of multi-modal sensors, the combination of events extracted from behavioural and physiological data were assessed for temporal accuracy and completeness. EEG event markers exhibited a low missing rate of 0.93% across 94,436 behavioural events. The timing precision of received EEG triggers was validated against corresponding behavioural events, with an offset of 0.09 ± 8.10 ms, equivalent to ±1 frame on a 144 Hz display. Webcam recordings were recorded in separate threads for each experimental block and saved as discrete MP4 files, enabling block-wise synchronisation and streamlined data handling.

## Usage Notes

The SENSE-42 dataset is organised in a flattened structure to facilitate ease of download and processing for individual or combined modalities. Data preprocessing and analysis pipeline were implemented using MNE^32^, NumPy^42^, Pandas ^43^, and SciPy ^44^. The source code for the experimental software and analysis pipeline is archived on Zenodo ^45^ (DOI: 10.5281/zenodo.20473861).

To reproduce the main experiment, the experimental software can be launched by opening explorer.psyexp in the experiment codebase archive ^45^ with PsychoPy v2024.2.3^30^, which manages stimulus presentation, timing, and event logging. For the analysis pipeline, environment setup and step-by-step usage instructions are provided on the archived Zenodo record^45^ and the corresponding GitHub repository (https://github.com/Catherine9811/HCI-SENSE-42/blob/master/README.md).

Designed around computer users, this dataset supports the investigation of tonic alertness dynamics during human-computer interaction. It enables the study of physiological fingerprints of behavioural performance and supports analysis of within- and between-subject factors in spontaneous user state fluctuations ^1;2^. Beyond user state analysis, the dataset allows for user behaviour analysis based on individual traits and graphical interface design, offering opportunities to explore the physiological and behavioural patterns related to the experience, habits, preferences of computer usage and cross-system design patterns. Furthermore, through the integration of physiological signals with interface interactions, this dataset may further support the development of physiological computing studies ^27^, where signals from the brain and body are transformed into adaptive inputs to optimise or personalise human-computer interfaces.

## Code Availability

The code used to generate, process and validate the dataset is archived on Zenodo ^45^ (DOI: 10.5281/zenodo.20473861) and is also available at the HCI-SENSE-42 repository (https://github.com/Catherine9811/HCI-SENSE-42) open sourced on GitHub.

## Acknowledgements

Sai Zhang was funded by the China Scholarship Council scholarship (CSC No. 202309210085). We thank Jingni Yan for providing template questions related to mind wandering and for sharing relevant research studies; Khadeejah Jahan for coordinating study support; and Chi-Hsu Wu, our lab manager, for maintaining the experimental room equipment and conditions.

## Author Contributions

Sai Zhang designed and conducted this research project and drafted the manuscript. Xinyu Bai provided support for code implementation and technical validation. Charles Hartley-O’Dwyer assisted with participant recruitment and study support. Hugh Warren assisted with the pilot study, participant recruitment and study support. Dr. Frederike Beyer supervised the study. Dr. Valdas Noreika conceptualised and supervised the study. All authors reviewed and approved the final version of the manuscript.

