## Supplementary figures and images for "A multimodal human-computer interaction dataset for neurocognitive user state evaluation"

### Supplementary Material S2

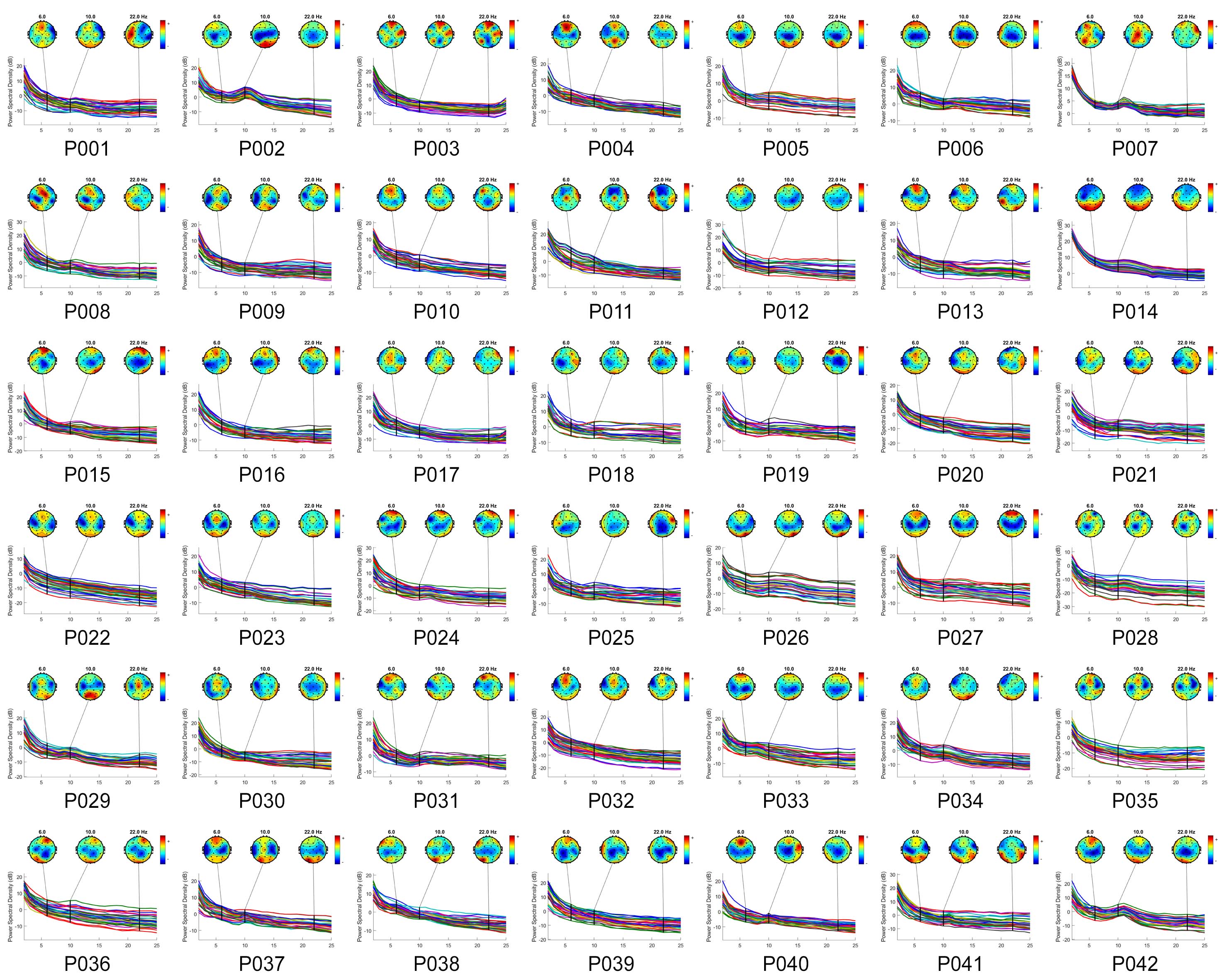
